# Akaike model selections of the vegetation structures and aerosphere factors in supporting lesser short nosed fruit bat (*Cynopterus brachyotis* Muller, 1838) populations in Asia mountainous paddy fields

**DOI:** 10.1101/2021.02.03.429676

**Authors:** Adi Basukriadi, Erwin Nurdin, Andri Wibowo, Jimi Gunawan

## Abstract

As an aerial and arboreal fauna, the abundances and populations of fruit bat *Cynopterus brachyotis* were influenced by the vegetation structures and aerosphere condition variables of fruit bat ecosystems. While mountaineous paddy field is an unique habitat since the trees are scarce and has exposure to the aerosphere variables including air temperature and humidity. Here this paper aims to select the best vegetation structures and aerosphere factors that support the abundance of *C. brachyotis* in mountainous paddy field landscape in West Java. The model selection was using AIC methodology by testing 15 models including 5 single models and 10 combination models of explanatory variables. Based on the model, tree height and combinations of tree height and elevation produced the best prediction for the bat abundances, as described by low values of AIC and the highest values of R^2^ and adjusted R^2^. For the best models, the AIC values ranged from 16.674 to 17.603, from 0.3404 to 0.4144 (R^2^), and 0.2461 to 0.2192 for adjusted R^2^. Regarding conservation of *C. brachyotis* and learning from the model, the conservation approaches mainly in mountainous paddy fields are encouraged to protect and conserve high altitude landscapes and trees with height > 10 m. Whereas the AIC results show lack of aerosphere variable effects on *C. brachyotis* (AIC: 19.346-20.406, R^2^: 0.1124-0.001353, and adjusted R^2^: −0.01444 − −0.1413).

## INTRODUCTION

Fruit bat (Pteropodidae) is one of the animals that is recognised as important in maintenaning healthy ecosystems and the provision of vital ecosystem services. Fruit bat has implication in the proliferation of at least 289 plant species and at least 186 of these plant species are having extrinsic human benefits ranging from timber, food to medicine. Fruit bat recently threatened with habitat conversion and destruction throughout Fruit bat distribution from subtropical to tropical Asia. In ecosystem, fruit bats require both terestrial and aerial suitable habitats. Especially for fruit bats and frugivores, the presences of trees mainly fruit trees are important. Trees are providing food resources mainly the fruiting trees and also shelter, protection and perching places for foraging fruit bats. This includes the lychee and mango trees with Mauritius fruit bat (*Pteropus niger*) (Tollington et al. 2019), *Pomophorus gambianus* and *Eidolon helvum* bats and the introduced neem tree (*Azadirachta indica*) (Ayensu 1974), and *Pteropus giganteus* and *Cynopterus sphinx* with Arecaceae (*Borassus flabellifer, Caryota urens*, *Washingtonia filifera*) and Annonaceae (*Polyalthiya longifolia*) for roosting habitats (Senthilkumar et al. 2012).

Bats are one of fauna groups that are dominating the aerosphere. The aerosphere, also known as a thin layer or atmosphere nearest the earth’s surface, is a dynamic habitat demanding posing distinct energetic and behavioural challenges and also providing opportunities to flying animals. Bats as flying animals living in this sphere should compromise specific aerosphere environmental conditions including wind, air temperature and humidity which may be largely decoupled from the below terrestrial habitat (O’Mara et al. 2019). Bats have advantage feature compared to other aerial organisms that occupy the aerosphere as bats extensively use this sphere for foraging, dispersal, migration, and behavioral interactions (Kalko et al. 2008). Differential use of the aerosphere is an important factor influencing bat assemblages, with species exhibiting distinct morphological, physiological, and sensory adaptations to different habitat types. While the aerosphere is not uniform, open-aerial species are highly successful at exploiting it. Bat individuals may potentially rely on visual or acoustic cues of others to move to higher and more productive altitudes, explore higher volumes in response to low foraging success. Bat also can monitor atmospheric conditions and change behaviour based on the potential aggregation of food patches at atmospheric boundaries.

Recently bat habitat in previously forest areas have been disturbed and changed into plantation and paddy fields. In mountaineous landscape, a montane forest has been converted into paddy field with remains of several trees. The conversion of closed habitat with dense trees to open habitat including paddy field may make bat experiencing more exposures on aerosphere variables including air temperature and humidity. In South East Asia, forest conversion to paddy field is common practice occurred in mountainous landscape. Whereas, how bat mainly lesser short nosed fruit bat *Cynopterus brachyotis* (Maryanto 1993, Suripto et al. 2006, Saridan 2016) that depends on the trees response to this condition is poorly understood. Then this paper aims to model the effects of vegetation and aerosphere variables on *C. brachyotis* population in mountainous landscape in West Java.

## MATERIALS AND METHODS

### Study area

This study was conducted in mountainous landscape located in Nagreg region in West Java Province, Indonesia. This landscape was dominated by mix of paddy fields, plantations, and settlements (Figure 1). The elevation was between 718 to 728 m above sea levels. In this study area, 3 transects with length of each transect was 1 km were located. The fruit bat, vegetation structure and aerosphere variable surveys were conducted in those transects in August 2020.

**Figure 1.**
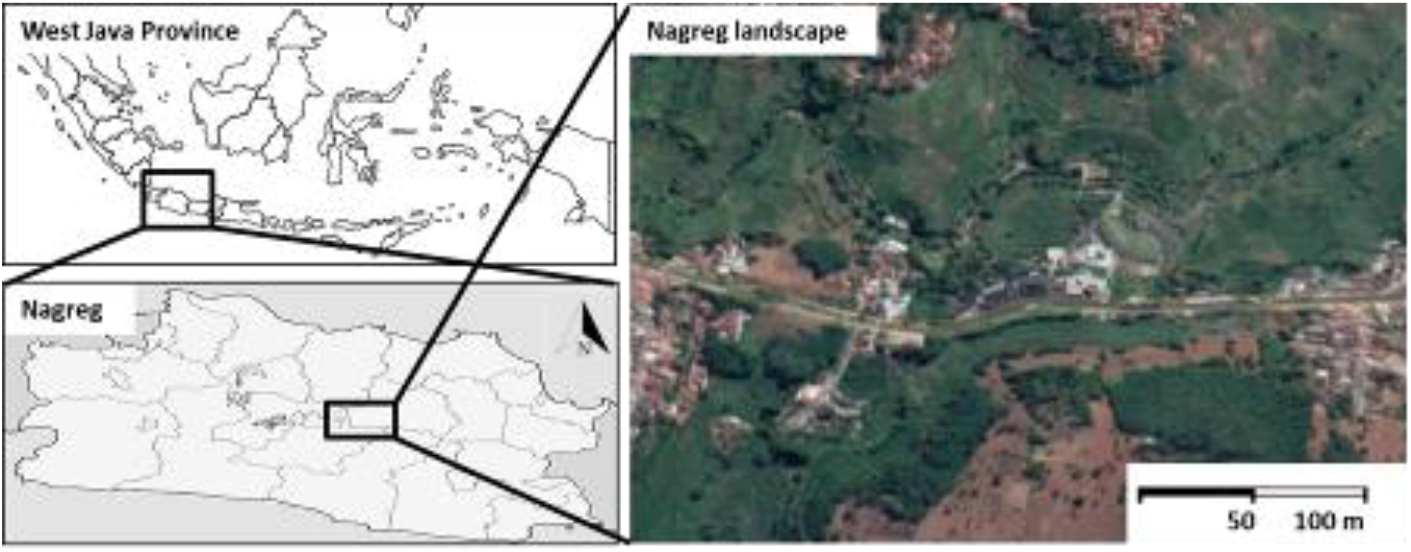
Location of study area in Nagreg landscape in West Java Province, Indonesia.

### Method

#### Vegetation and habitat surveys

Vegetation surveys were conducted in August 2020 following the fruit bat surveys from 17.00 to 19.00. The vegetations that where passed and perched by bats were recorded. The recorded variables including size of canopy cover and tree height. The surveys were conducted within a square plot sizing 10 m x 10 m standardized for tree surveys. Only vegetation falls into tree category was recorded and excluded shrub, herb, and understorey vegetations. The measured aerosphere variables measured during the surveys included the elevation, air temperature and humidity. As an aerial fauna that spend most of the activities in air, the aerosphere variables can affect the bat abundances.

#### Cynopterus brachyotis surveys

*C. brachyotis* abundance observations were conducted following methods as described by several authors. The observations were conducted at noon from 17.00 to 19.00 following *C. brachyotis* activity times within 10 m x 10 m grid in each sampling location. *C. brachyotis* abundances were recorded and denoted as individuals/100 m^2^.

### Data analysis

Vegetation structures and aerosphere variables model were developed using Akaike Information Criterion (AIC) following Duff & Morrell (2007), Ingersoll (2010) and Hoffman et al. (2019) to evaluate and assemble models of vegetation structure and *C. brachyotis* abundance. The AIC was developed using the linear regression. The measured parameters included in AIC are R^2^ and adjusted R^2^. To build the model, 5 explanatory variables including tree covers in %/100 m^2^, tree heights in m, air temperature in °C, air humidity in %, and elevations in m with 10 combinations of those variables were included in the analysis to develop the model.

## RESULTS AND DISCUSSION

### Akaike model selection

Among all 5 explanatory variables and 10 combinations of those variables, tree height and combinations of tree height and elevation produced the best prediction for the abundances, as described by low values of AIC and the highest values of R^2^ and adjusted R^2^ (Table 1). For the best models, the AIC values ranged from 16.674 to 17.603, 0.3404 to 0.4144 (R^2^), and 0.2461 to 0.2192 for adjusted R^2^ (Table 1). Whereas the AIC results show lack of aerosphere variable effects on *C. brachyotis* (AIC: 19.346-20.406, R^2^: 0.1124-0.001353, and adjusted R^2^: −0.01444 − −0.1413).The correlation of *C. brachyotis* with vegetation structure can be observed in tree height and cover (Figure 2). While weak correlations were observed between *C. brachyotis* abundance and aerosphere variables including air temperature and humidity (Figure 2). The PCA analysis (Figure 3) also confirms the influences of tree height on bat abundances that more significant than air temperature and humidity factors.

**Figure 2.**
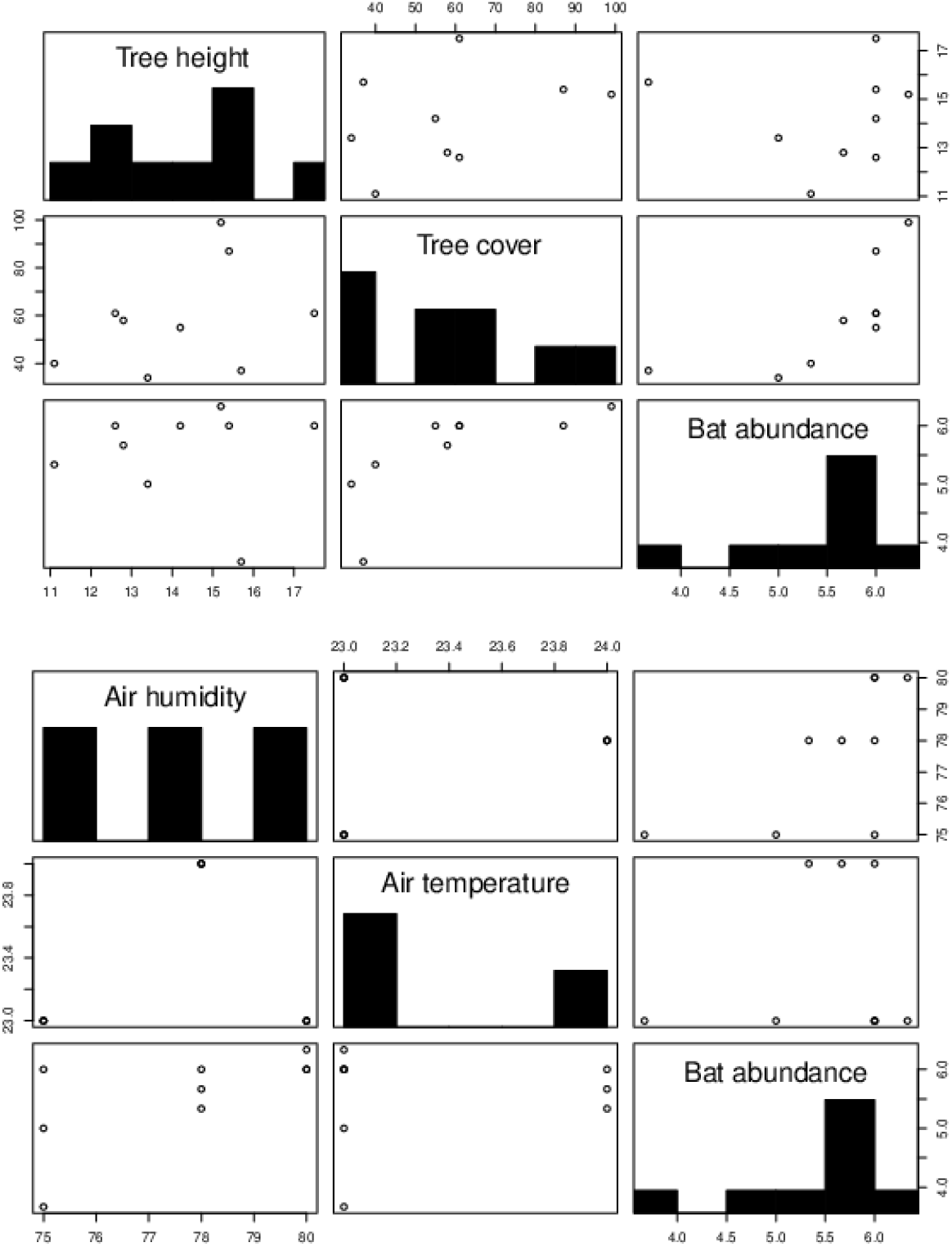
Correlation plots of fruit bat *Cynopterus brachyotis* abundances ((inds./100 m^2^) with vegetation structure variables (tree covers in %/100 m^2^ and tree heights in m) and aerosphere variables (air temperature in °C and humidity in %).

**Figure 3.**
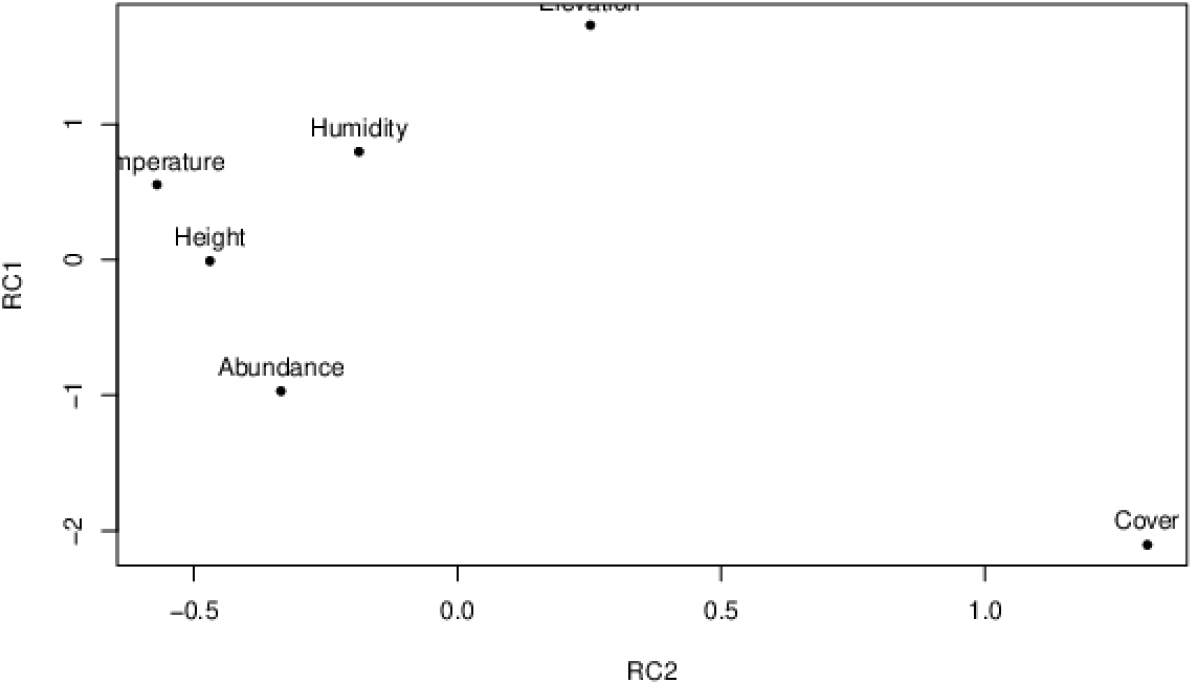
PCA plots of fruit bat *Cynopterus brachyotis* abundances (inds./100 m^2^) with vegetation structure variables (tree covers in %/100 m^2^ and tree heights in m) and aerosphere variables (air temperature in °C and humidity in %).

**Table 1.** Akaike Information Criteria (AIC), R^2^, adjusted R^2^ for the multiple regression models between abundance and vegetation (tree covers, heights) and aerosphere variables (air temperature, humidity), combination of vegetation and aerosphere variables. The values with asterisk signs show the best models selected.

| Model | AIC | R<sup>2</sup> | Adj R<sup>2</sup> | F | P |
| --- | --- | --- | --- | --- | --- |
| Tree cover | 19.658 | 0.08106, | -0.05021 | 0.6175 | 0.4577 |
| Tree height | 16.674* | 0.3404 | 0.2461 | 3.612 | 0.09914 |
| Elevation | 19.995 | 0.05018 | -0.0855 | 0.3698 | 0.5623 |
| Air temperature | 19.346 | 0.1124 | -0.01444 | 0.8861 | 0.3779 |
| Air humidity | 20.406 | 0.001353 | -0.1413 | 0.009482 | 0.9252 |
| Tree cover + height | 18.589 | 0.3466 | 0.1287 | 1.591 | 0.279 |
| Tree cover + elevation | 20.676 | 0.176 | -0.09862 | 0.6409 | 0.5594 |
| Tree cover + air temperature | 20.849 | 0.1607 | -0.1191 | 0.5742 | 0.5913 |
| Tree cover + air humidity | 21.181 | 0.1285 | -0.162 | 0.4424 | 0.6619 |
| Tree height + elevation | 17.603* | 0.4144 | 0.2192 | 2.123 | 0.2008 |
| Tree height + air temperature | 18.104 | 0.3808 | 0.1744 | 1.845 | 0.2374 |
| Tree height + humidity | 18.266 | 0.3696 | 0.1595 | 1.759 | 0.2505 |
| Elevation + air temperature | 21.288 | 0.1181 | -0.1759 | 0.4017 | 0.6859 |
| Elevation + air humidity | 21.288 | 0.1181 | -0.1759 | 0.4017 | 0.6859 |
| Air temperature + air humidity | 21.288 | 0.1181 | -0.1759 | 0.4017 | 0.6859 |

## Discussion

Akaike model selection has been widely used in biology in general and ecology studies in particular. This approach is very useful since the biological and ecological process were not a standalone process, whereas it was the combinations of several explanatory variables. Akaike model selection methods are very applicable in many fields. Humagain et al. (2017) have used this approach to deal with the complexity in forest management using remotes sensing since different vegetation types have different spectral signatures. By using Akaike in their research they have derived models with multiple predictor variables for each vegetation type rather than using a single model. Based on the AIC model, they found that for the management of conifer-dominated forest, should use Landsat bands 2, 3, 4, and 7 along with simple ratios, normalized vegetation indices, and the difference vegetation index. While individual Landsat bands 4 and 2, simple ratio, and normalized vegetation index are the best for the management of aspen-dominated areas.

The AIC alone has been widely used in bat studies. The aim of AIC is to assess and model which environmental factor is required and the impact of those factors mainly in influencing the bat populations. In northern hemisphere, bat populations require snags for roosting sites as observed in many bats inhabiting coniferous forests. By using AIC analysis, Lacki et al. (2012) have demonstrated that the snag condition model was to be most parsimonious in all for roosting sites. The influence of numerous environmental variables including anthropogenic factors using AIC can also be seen in research by Kerbiriou et al. (2015) in assessing the bat population roosting sites inside the railway tunnel. According to AIC values, they found that the most important variables affecting bat population inside tunnel were appeared to be the yearly trend, followed by presence of railway traffic, the group of winter weather variables, and secondarily the group of summer variables.

In our study and Akaike model shows that the aerosphere factors have limited effects on the fruit bat populations. This result is comparable to O’Mara et al. (2019). Based on their study, there was no explanatory relationship between the bat presences with wind speed, air temperature, wind direction and air pressure. In Nagreg landscape and based on AIC values, air temperature has high AIC value. In comparison between observed aerosphere variables, AIC value of air temperature was lower than humidity and indicated that temperature has more effects on bat populations. Scanlon and Petit (2008) have reported an increase in bat activity affected by air temperature.

In our model, the elevation above sea level combined with the tree height is the best model to depict the fruit bat population. The elevation is considered as one of factors influencing bat populations. Cryan et al. (2000) have observed positive relationship between male distribution and elevation whereas female bats have inverse relationship with elevation factors. In our AIC model, elevation is not a standalone factor that affects the fruit bat population. Instead of single factor, the elevation should be combined with the tree height variable to provide significant impacts on fruit bat population. The less effect of standalone elevation variable on fruit bat population in our study is in agreement with Georgiakakis et al. (2010).

The conservation of bat in Asia is challenged by availability of information on which environment factors that important for bat that remain poorly understood. To conclude this paper has provided a robust method along with its empirical evidence to support land use planning and management efforts in ensuring and supporting fruit bat populations.

## Abbreviations

AIC: (Akaike Information Criterion)

